# Enrofloxacin shifts intestinal microbiota and metabolic profiling and hinders recovery from *Salmonella enterica* subsp. *enterica* serovar *Typhimurium* infection in neonatal chickens

**DOI:** 10.1101/2020.07.13.201830

**Authors:** Boheng Ma, Xueran Mei, Changwei Lei, Cui Li, Yufeng Gao, Linghan Kong, Xiwen Zhai, Hongning Wang

## Abstract

Enrofloxacin is an important antibiotic used for prevention and treatment of *Salmonella* infection in poultry in many countries. However, oral administration of enrofloxacin may lead to the alterations in the microbiota and metabolome in the chicks intestine, thereby reducing colonization resistance to the *Salmonella* infection. To study the effects of enrofloxacin on chicken cecal *Salmonella*, we used different concentrations of enrofloxacin to feed 1-day-old chickens, followed by oral challenge with *Salmonella enterica* subsp. *enterica* serovar *Typhimurium* (*S. Typhimurium*). We then explored the distribution patterns of *S. Typhimurium in vivo* in intestinal contents using quantitative polymerase chain reaction and microbial 16S amplicon sequencing on days 7, 14, and 21. Metabolome sequencing was used to explore the gut metabolome on day 14. Faecalibacterium and Anaerostipes, which are closely related to the chicken intestinal metabolome, were screened using a multi-omics technique. The abundance of *S. Typhimurium* was significantly higher in the enrofloxacin-treated group than in the untreated group, and *S. Typhimurium* persisted longer. Moreover, the cecal colony structures of the three groups exhibited different characteristics, with *Lactobacillus* reaching its highest abundance on day 21. Notably, *S. Typhimurium* infection is known to affect the fecal metabolome of chickens differently. Thus, our results suggested that enrofloxacin and *Salmonella* infections completely altered the intestinal metabolism of chickens.

**IMPORTANCE:** In this study, we examined the effects of *S. Typhimurium* infection and enrofloxacin treatment on the microbial flora and metabolite synthesis in chicken guts in order to identify target metabolites that may cause *S. Typhimurium* colonization and severe inflammation and to evaluate the important flora that may be associated with these metabolites. Our findings may facilitate the use of antibiotics to prevent *S. Typhimurium* infection.

## INTRODUCTION

Foodborne diseases are associated with high morbidity and mortality rates worldwide and pose major challenges to food safety and economics (1). *Salmonella* is an important foodborne pathogen of the family Enterobacteriaceae. This pathogen causes millions of infections worldwide each year, both in humans and livestock. *Salmonella* has been shown to cause intestinal inflammation and barrier dysfunction in chickens, thereby affecting chicken performance (2). Currently, more than 2,600 serotypes of *Salmonella* are known, with varying pathogenicity and host specificity. *Salmonella enterica* subsp. *enterica* serovar *Typhimurium* (*S. Typhimurium*) is a typical representative of non-host-specific *Salmonella* found in poultry.

The main route of infection with *Salmonella* in poultry is the fecal-oral route. For multiserotype infections, bacteria mainly colonize in the ceca of poultry. After colonizing the intestines, these bacteria invade the intestinal epithelial cells and dendritic cells and reach the submucosa to be phagocytosed by macrophages. The bacteria then replicate in and colonize the spleen and liver, causing systemic infections mediated by bacterium/macrophage interactions (3,4). In mice infected with *Salmonella*, macrophages produce IL-12 and IL-18 and promote IFN-γ secretion by natural killer (NK) cells (5).

Owing to the widespread use of antibiotics in agriculture and animal husbandry, *Salmonella* resistance has rapidly increased. In 2013, China alone consumed approximately 927,000 tons of antibiotics (6). Moreover, the consumption of antibiotics as veterinary drugs increased from 46% to 52% between 2007 and 2013. Enrofloxacin, florfenicol, lincomycin, and amoxicillin are among the most commonly consumed veterinary antibiotics in China (7). Enrofloxacin is the main antibiotic used for treating *Salmonella* infection in poultry in many countries, and the concentration of enrofloxacin in chicken manure in China was found to be 1,421 mg/kg (8).

Quinolones, such as enrofloxacin, affect the composition and number of bacterial species in the chicken gut (9), thereby altering resistance to pathogen invasion. Indeed, bacteria residing in the gut can mediate the colonization of *S. Typhimurium* in the body by producing short-chain fatty acids (SCFAs), such as propionic acid, which inhibits *S. Typhimurium* growth by influencing the intracellular pH (10). By producing SCFAs, lactic acid bacteria inhibit *Salmonella* colonization in the colon and jejunum, accelerate *Salmonella* clearance from stools, and reduce *Salmonella* transfer from the small intestine to the spleen (11). The concentrations of SCFAs and butyric acid are inversely proportional to the degree of inflammation (12).

Some Enterobacteriaceae in the normal gut flora can mediate *Salmonella* multiplication through competition for oxygen (13). Additionally, antibiotics reduce the number of butyrate-producing *Clostridium perfringens* in the intestinal flora, shifting the host cells from oxidative metabolism to lactic acid fermentation and increasing the level of lactic acid in the intestine; *Salmonella* then gain a colonization advantage by using lactic acid (14). Intestinal metabolism is also affected by the gut flora composition. Using antibiotics increases fucose and sialic acid contents in the intestine, thereby affecting *Salmonella* and *C. difficile* colonization (15,16). Similarly, antibiotics also decrease the levels of secondary bile acids in the intestine and inhibit *C. difficile* colonization (16).

Accordingly, in this study, we investigated the effects of enrofloxacin on *Salmonella* colonization in chickens and the relationship between intestinal flora and *Salmonella* colonization. We also evaluated changes in microbial metabolites in the ceca of enrofloxacin-treated chickens infected with *Salmonella* and used multiomics association analyses to determine the correlations between microbial community structure and microbial metabolites.

## RESULTS

### Response of *Salmonella* to enrofloxacin

First, *S. Typhimurium* abundance in the samples was evaluated. *S. Typhimurium* was not isolated from C1, C2, or C3 groups. On day 7, the highest abundance of *S. Typhimurium* was found in the cecum contents of group E1 (6.128 log_10_CFU/g), followed by that of groups E3 and (5.090 and 3.945 log_10_CFU/g, respectively; *P* < 0.001). The abundances of *S. Typhimurium* in the heart, liver, and spleen of group E1 were significantly higher than those in the other two groups (*P* < 0.001). On day 14, the abundance of *S. Typhimurium* in group E1 (3.371 log_10_CFU/g) was significantly lower than those in groups E2 (4.829 log_10_CFU/g) and E3 (4.366 log_10_CFU/g; *P* < 0.01). Moreover, the abundances of *S. Typhimurium* were significantly lower in heart, liver, and spleen samples from group E1 than those from groups E2 and E3, except for heart samples from group E3. On day 21, the relative abundances of *S. Typhimurium* were 3.170, 5.146, and 5.636 log_10_CFU/g in groups E1, E2, and E3, respectively; that in group E1 was significantly lower than those in the other two groups. However, no *S. Typhimurium* was detected in the heart, liver, or spleen samples of group E1 (Figure 1).

**Figure 1.**
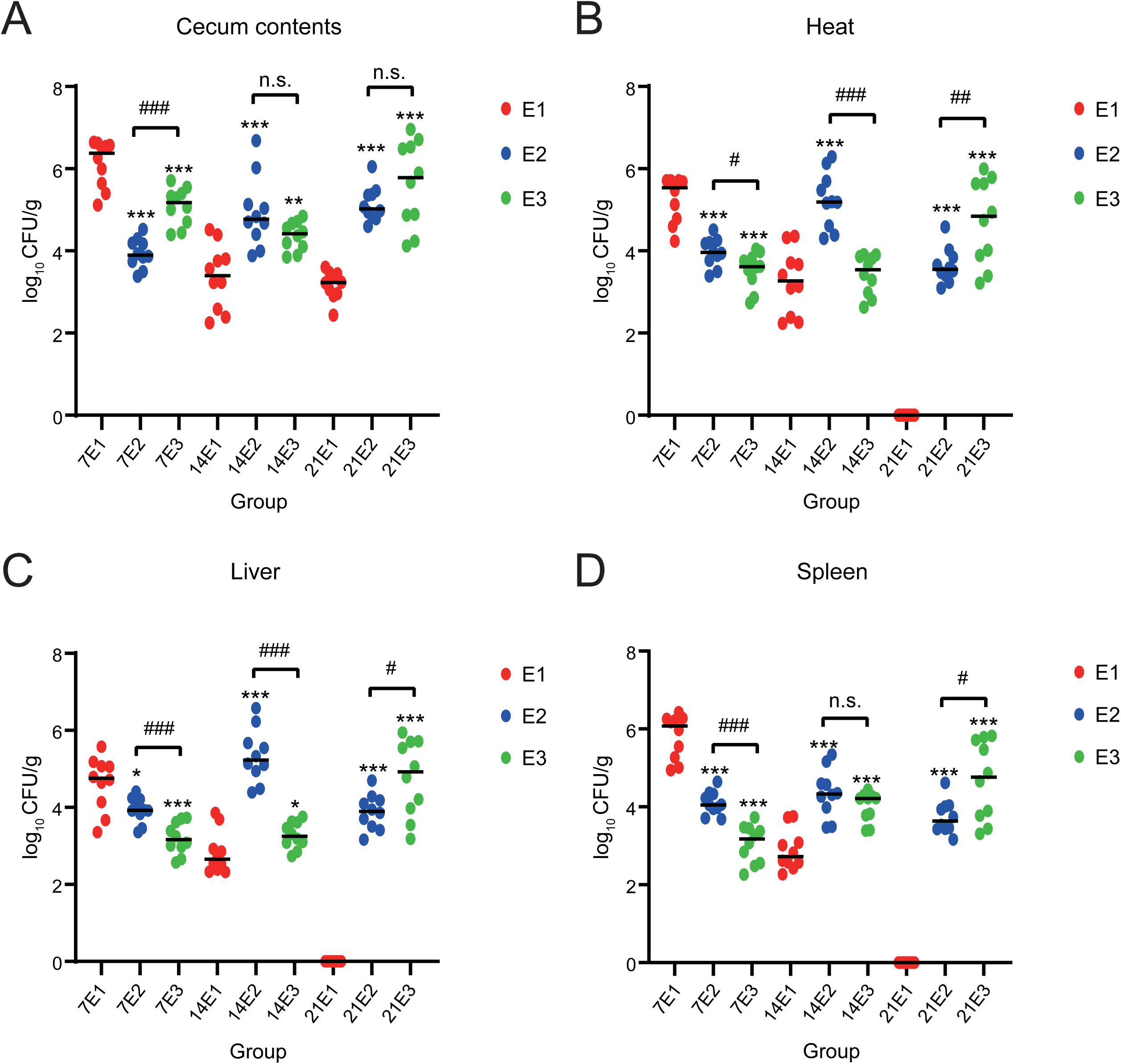
*Salmonella* abundance. (A) Cecum contents, (B) heart, (C) liver, and (D) spleen. The *x*-axis represents log_10_ copies/g, and the *y*-axis represents different groupings. Compared with the same sampling point in group E1 group, *P* < 0.001, indicated by ***; 0.001 ≤ *P* ≤ 0.01, indicated by **; and 0.01 < *P* ≤ 0.05, indicated by *. *P* > 0.05, no mark. Comparing groups E2 and E3, *P* < 0.001, indicated by ###; 0.001 ≤ *P* ≤ 0.01, indicated by ##; and 0.01 < *P* ≤ 0.05, indicated by #. *P* > 0.05, n.s.

### Expression levels of immune-related genes following enrofloxacin treatment

On day 14, *MUC*2 expression was significantly lower in group E3 than in group E1 (*P* < 0.05), *TLR*21 expression was significantly higher in group E3 than in groups E1 and E2 (*P* < 0.05), *IL-* 1*β* and *LITAF* levels were significantly higher in group E3 than in group E1 (*P* < 0.05), *IL-*18 and *OCLM* levels were significantly higher in group E1 than in groups E2 and E3 (*P* < 0.05), and *TLR*4 expression did not differ significantly among the three groups (Figure 2).

**Figure 2.**
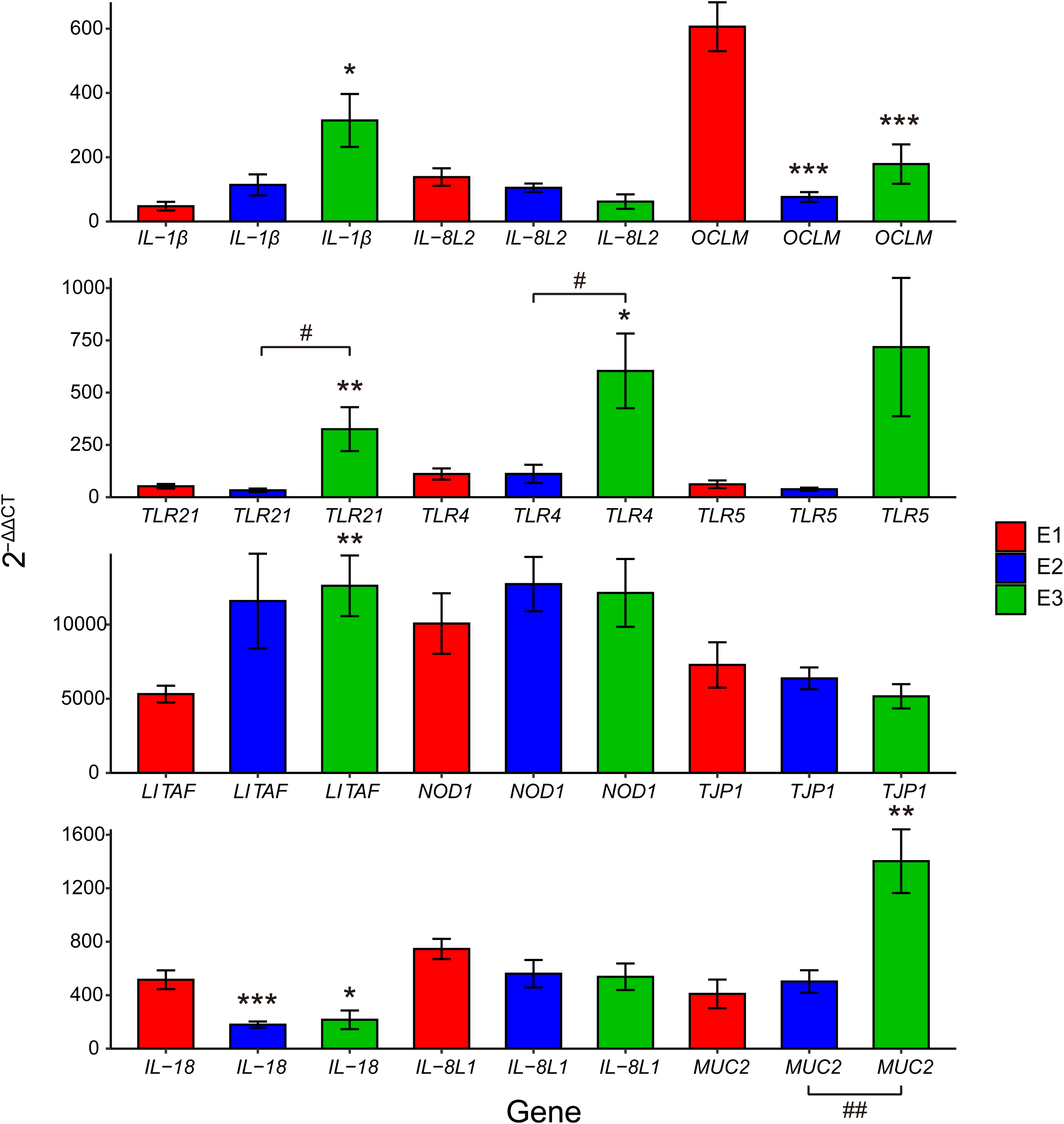
Gene expression. The β-actin gene was used as an endogene, and relative expression was calculated by normalization to this gene using the 2^−ΔΔCT^ method. (A) *IL-1β*, *IL-8L2*, and *OCLM*; (B) *TLR4*, *TLR5*, and *TLR21*; (C) *LITAF*, *NOD*, and *TJP*; (D) *IL-18*, *IL-8L1*, and *MUC2* on the *x*-axis, with different genes on the *y*-axis. Compared with group E1 at the same sampling point, *P* < 0.001, indicated by ***; 0.001 ≤ *P* ≤ 0.01, indicated by **; and 0.01 < *P* ≤ 0.05, indicated by *. *P* > 0.05, no mark. Comparing groups E2 and E3, *P* < 0.001, indicated by ###; 0.001 ≤ *P* ≤ 0.01, indicated by ##; and 0.01 < *P* ≤ 0.05, indicated by #. *P* > 0.05, n. s.

### Effects of enrofloxacin on the microbial composition and structure

By 16S rRNA sequencing of cecum colonies, as an increased number of samples was sequenced, the dilution curve flattened, indicating that the sequencing data were saturated and that the sequencing results could reflect the majority of the samples (Figure S1). On day 7, significant differences were observed in the microbial diversity of the ceca among groups E1, E2, and E3 (*P* < 0.05). Group E1 had the lowest diversity, whereas group E3 had the highest diversity. On day 14, the microbial diversity of all experimental groups was higher than that on day 7 (*P* < 0.05). However, on day 14, no differences were observed in the Good’s coverage index, ACE index, observed species index, or Chao index between the groups (*P* > 0.05). However, Shannon’s and Simpson’s diversity indices were significantly different (*P* < 0.05). On day 21, microbial diversity was higher in group E2 than in groups E1 and E3 (*P* < 0.05; Figure S2).

On day 7, the microbial community structures of cecum samples from groups E1, E2, and E3 differed significantly in the hierarchical clustering tree. The differences in the microbial community structure of the samples collected on days 14 and 21 gradually decreased, and the samples were mixed together in the clustering tree (Figure 3). PCA showed that on day 7, the structure of the cecum flora was isolated in groups E1, E2, and E3 owing to the use of different concentrations of antibiotics. However, on days 14 and 21, the antibiotic concentrations had no significant effect on the structure of the cecum flora (Figure S3).

**Figure 3.**
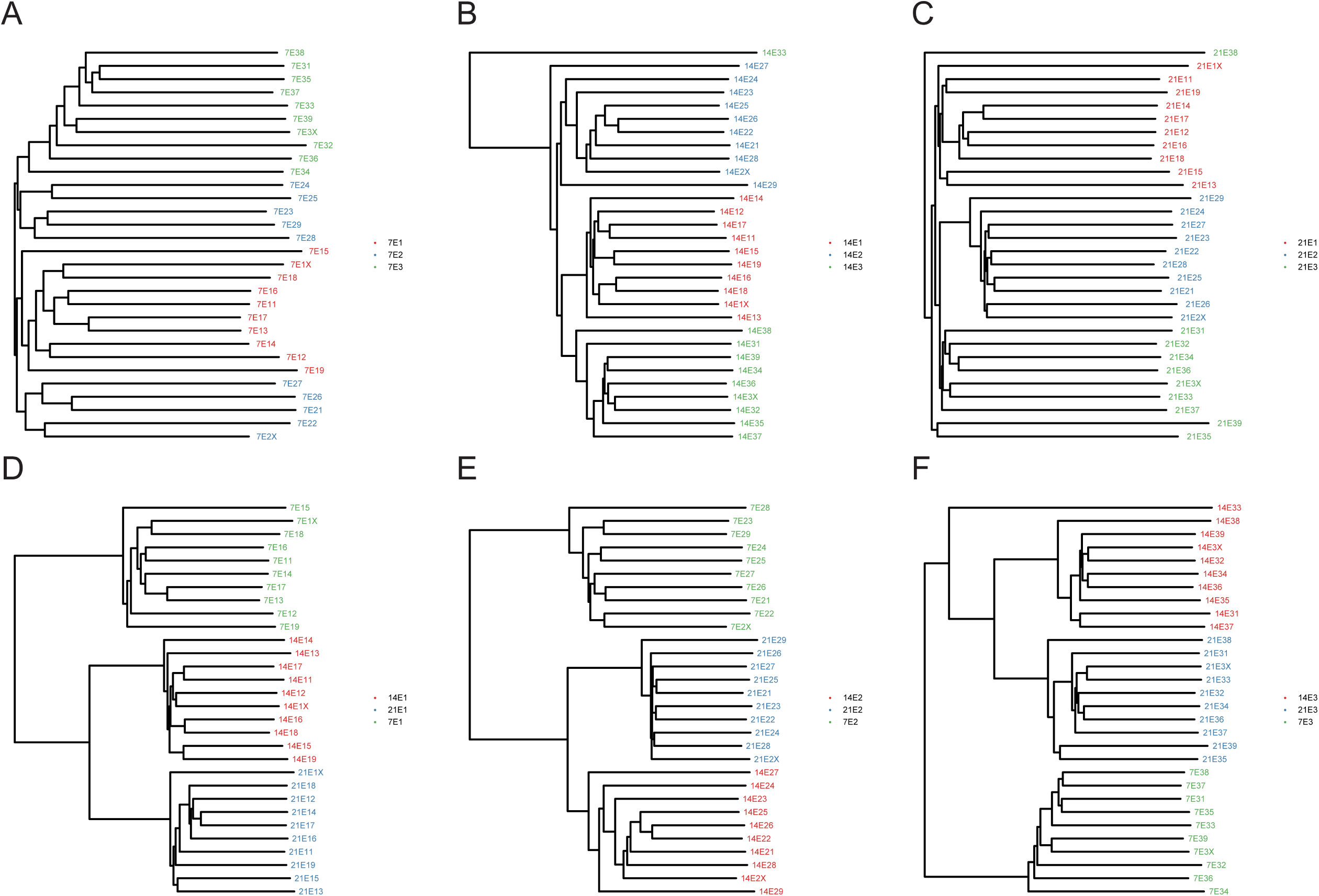
Beta-diversity analysis of the cecum microbiota after *Salmonella* infection. According to the statistical results of the variability of each sample, a clustering analysis was performed on the samples in groups E1, E2, and E3 at the three sampling points and the intersample distance was calculated. (A, B, C) Cluster trees for groups E1, E2, and E3 on days 7, 14, and 21. (D, E, F) Cluster trees for groups E1, E2, and E3 at the three sampling points.

An UpSetR chart shows the common and unique OTUs of different categories of chicken cecum microbes. On day 7, 193 OTUs from the three groups were found to be common, whereas 22, 24, and 55 OTUs from groups E1, E2, and E3, respectively, were found to be unique. On day 14, 391 OTUs from the three groups were found to be shared, whereas 10, 34, and 22 OTUs from groups E1, E2, and E3, respectively, were found to be unique. On day 21, 396 OTUs from the three groups were found to be shared, whereas 15, 44, and 18 OTUs from groups E1, E2, and E3, respectively, were found to be unique. This showed that as the chicken intestinal flora matured, the OTUs shared among different groups increased, unique OTUs decreased, and the microbial community structures tended to be similar. In total, 200 OTUs were found to be shared by group E1 at the three sampling points. On days 7, 14, and 21, the numbers of unique OTUs were 19, 56, and 57, respectively. In total, 218 OTUs were found to be shared by group E2 at the three sampling points. On days 7, 14, and 21, the numbers of unique OTUs were 9, 50, and 80, respectively. In total, 229 OTUs were found to be shared by group E3 at the three sampling points. On days 7, 14, and 21, the numbers of unique OTUs were 18, 40, and 74, respectively. Thus, with maturation of intestinal microbes, the microbial OTUs in the cecum became more and more abundant (Figure 4).

**Figure 4.**
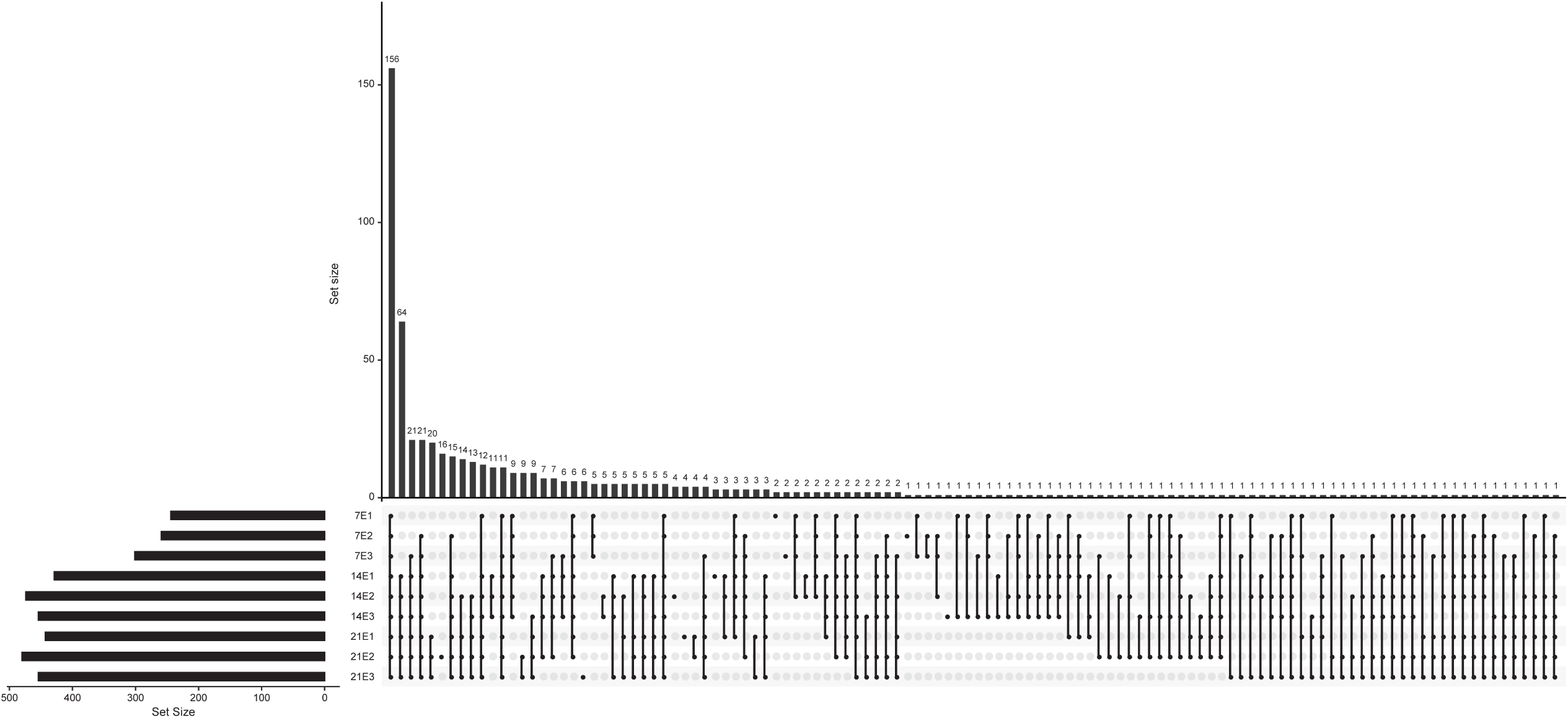
The left histogram shows the number of OTUs on the x-axis and the name of the comparison group on the y-axis. The upper right histogram shows the intersection of the different comparison groups on the x-axis and the number of OTUs on the y-axis. Each column in the lower right indicates the relationship between the left comparison group and the intersection above. Dark dots indicate that the intersection contains the comparison group, light dots indicate that the intersection does not contain the comparison group, and dark dots are connected by lines.

We also measured the relative abundance of the cecal flora at the phylum (Figure 5A) and genus levels (Figure 5B). At the phylum level, three dominant bacterial groups with a relative abundance of more than 1% were identified in the experimental groups (i.e., Firmicutes, Bacteroidetes, and Proteobacteria). Among these, the abundance of Firmicutes was highest in groups E1 and E2 on day 7 and in group E3 on day 14. On day 14, the abundance of Bacteroidetes was higher in group E2 than in the other two groups. On day 21, the abundance of Bacteroidetes reached its highest level in the three groups. Moreover, the abundance of Proteobacteria was highest in group E1 on day 14 and in the other two groups on day 7.

**Figure 5.**
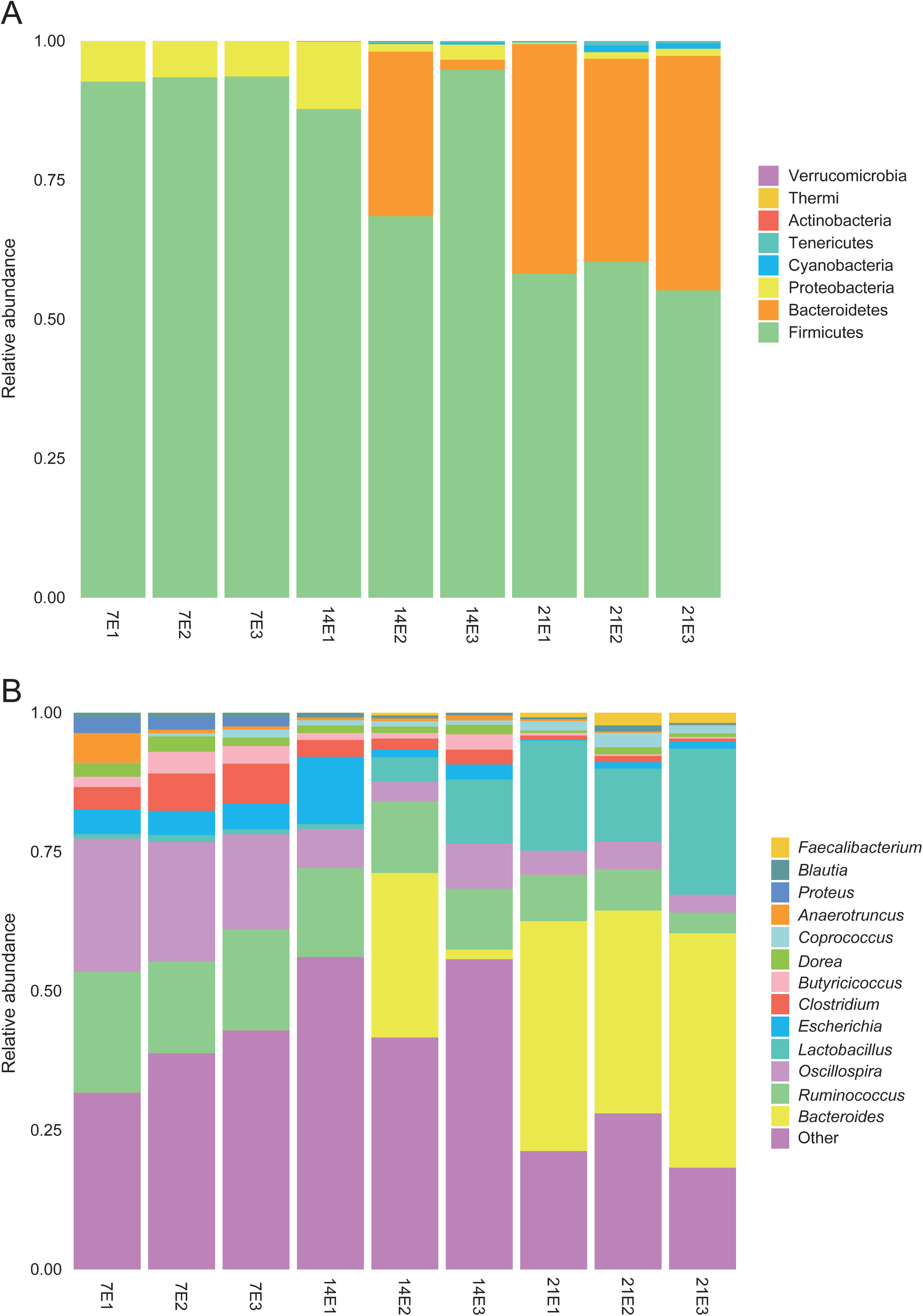
Average relative abundance of microbial species in the cecum at the (A) phylum level and (B) genus level.

At the genus level, the top 10 most abundant microbes in each of the three groups included *Coprobacillus*, *Coprococcus*, *Blautia*, *Lactobacillus*, *Anaerotruncus*, *Proteus*, *Dorea*, *Butyricicoccus*, *Escherichia*, *Clostridium*, *Ruminococcus*, *Oscillospira*, *Bacteroides*, and *Faecalibacterium*. On day 7, the relative abundances of *Dorea* and *Anaerotruncus* were significantly higher in group E1 than in the other two groups, and the relative abundances of *Clostridium* and *Coprococcus* were significantly higher in group E3 than in groups E1 and E2. On day 14, no significant differences were observed in the relative abundances of the 10 dominant bacterial groups. However, the relative abundances of *Ruminococcus*, *Clostridium*, and *Bacillus* were highest in group E1, and those of *Oscillospira*, *Clostridium*, *Butyricicoccus*, *Escherichia*, *Lactobacillus*, and *Brucella* were lowest in the three groups. In group E3, the relative abundances of *Ruminococcus* and *Coprobacillus* were lowest among the three groups, whereas the relative abundances of *Oscillospira*, *Dorea*, *Butyricicoccus*, *Escherichia*, *Anaerotruncus*, and *Lactobacillus* were highest. On day 21, the relative abundance of *Ruminococcus* was significantly higher in group E1 than in the other two groups. On day 21, the relative abundances of *Clostridium*, *Dorea*, and *Brucella* were significantly higher in group E2 than in the other groups. Additionally, the relative abundances of *Oscillospira*, *Clostridium*, *Dorea*, *Escherichia*, and *Anaerotruncus* were highest in group E1 on day 7, and those of *Coprococcus* and *Lactobacillus* were highest in group E1 on day 21. In group E2, the relative abundances of *Ruminococcus*, *Oscillospira*, *Clostridium*, *Dorea*, *Butyricicoccus*, and *Escherichia* were highest on day 7, and those of *Coprococcus*, *Lactobacillus*, and *Faecalibacterium* were highest on day 21. In group E3, the relative abundances of *Ruminococcus*, *Oscillospira*, *Clostridium*, *Butyricicoccus*, and *Coprobacillus* were highest on day 7, that of *Dorea* was highest on day 14, and that of *Lactobacillus* was highest on day 21 (Figure 6). On day 21, the relative abundances of the three groups of lactobacilli peaked, potentially because the intestinal tracts of infected chickens were conducive to the growth of lactic acid bacteria.

**Figure 6.**
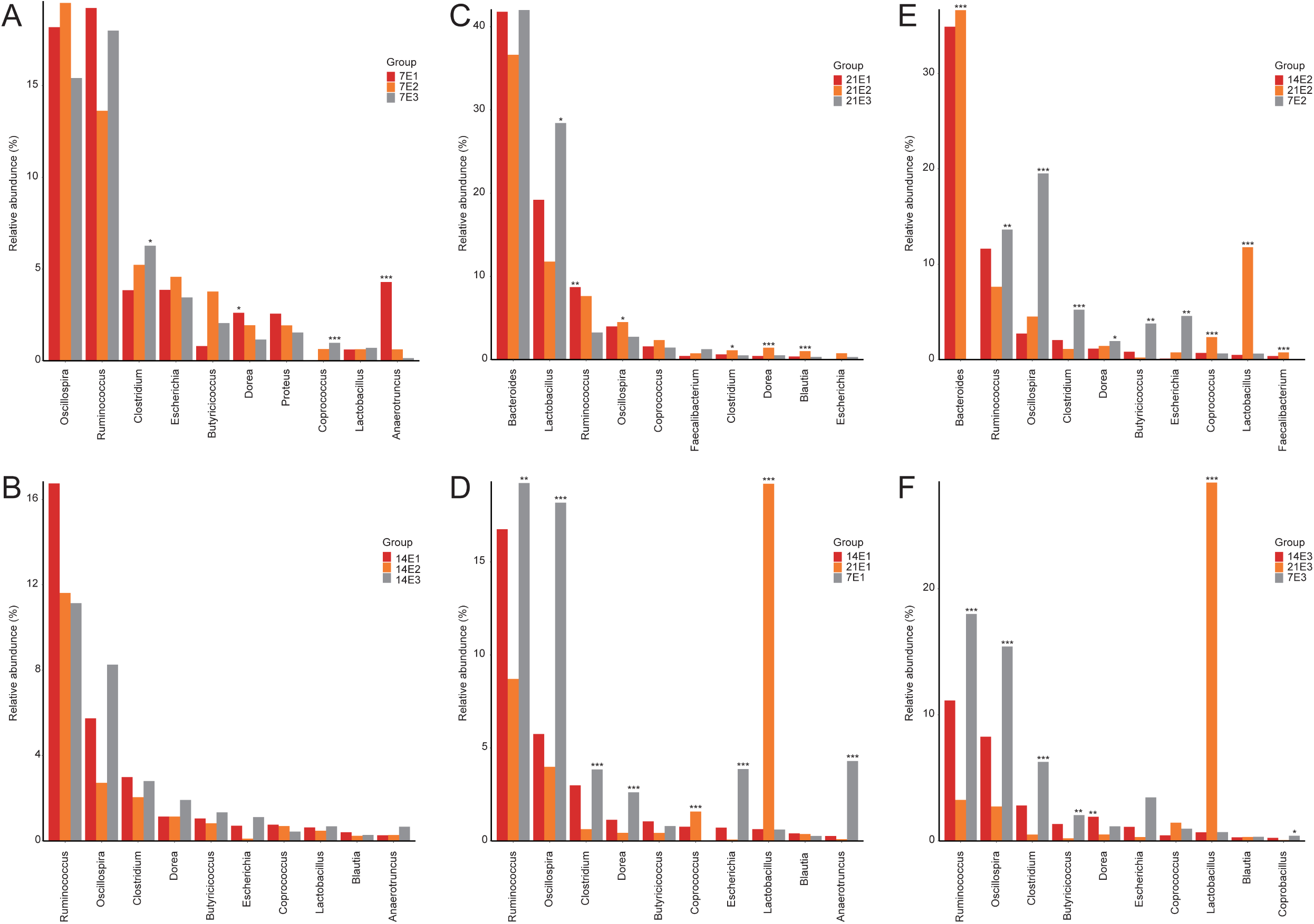
Key species difference comparisons, selecting the top 10 species with the highest abundance, demonstrating the mean relative abundance of each group and the significance of the difference test. (A, B, C) Groups E1, E2, and E3 groups with the top 10 species with the highest abundance on days 7, 14, and 21. (D, E, F) Alpha diversity of groups E1, E2, and E3 at the three sampling points. *P* < 0.001, denoted by **; 0.001 ≤ *P* ≤ 0.01, denoted by **; and 0.01 < *P* ≤ 0.05, denoted by *. P > 0.05, no mark.

LefSe analysis of the bacterial species composition in groups E1, E2, and E3 at the three sampling time points revealed that on day 7, *Coprococcus* and *Clostridium* were the most important bacteria differentiating group E1 from the other two groups, with the characteristic colony in group E2 being *Faecalibacterium*. *Salmonella*, *Streptococcus*, and *Anaerotruncus* showed the most obvious differences between the E3 group and the other two groups. *Salmonella* could easily distinguish group E3 from the other groups. On day 14, Ruminococcaceae, Clostridia, and Clostridiales in group E1 showed the most significant differences between the three groups, and *Faecalibacterium*, Aerococcaceae, Bacteroidetes, Bacteroidia, Bacteroidales, *Bacteroides*, and Bacteroidaceae in group E2 showed the most significant differences among the three groups. In group E3. Firmicutes, Oscillospira, Dehalobacteriaceae. Dehalobacterium, Anaerostipes. Erysipelotrichales, Erysipelotrichaceae, and Erysipelotrichi showed the most significant differences among the groups. On day 21, Microbacteriaceae, *Leucobacter*, and *Ruminococcus* in group E1 showed the most significant differences among the three groups. Moreover, *Defluviitalea*, *Coprobacillus*, Eubacteriaceae, *Anaerofustis*, Leuconostocaceae, *Weissella*, *cc_*115, Erysipelotrichales, Erysipelotrichi, *Erysipelptrichaceae*, *Dehalobacteriaceae*, *Dehalobacterium*, *Eubacterium*, *Clostridium*, *Blautia*, *Dorea*, *Coprococcus*, *Oscillospira*, *Epulopiscium*, Lachnospiraceae, Ruminococcaceae, Clostridia, and Clostridiales showed the most significant differences in group E2. In group E3, *Staphylococcus*, Campylobacterales, Helicobacteraceae, *Akkermansia*, Verrucomicrobiaceae, Verrucomicrobiales, *Bacillus*, Lactobacillales, *Lactobacillus*, and Lactobacillaceae showed the most significant differences among the three groups (Figure 7).

**Figure 7.**
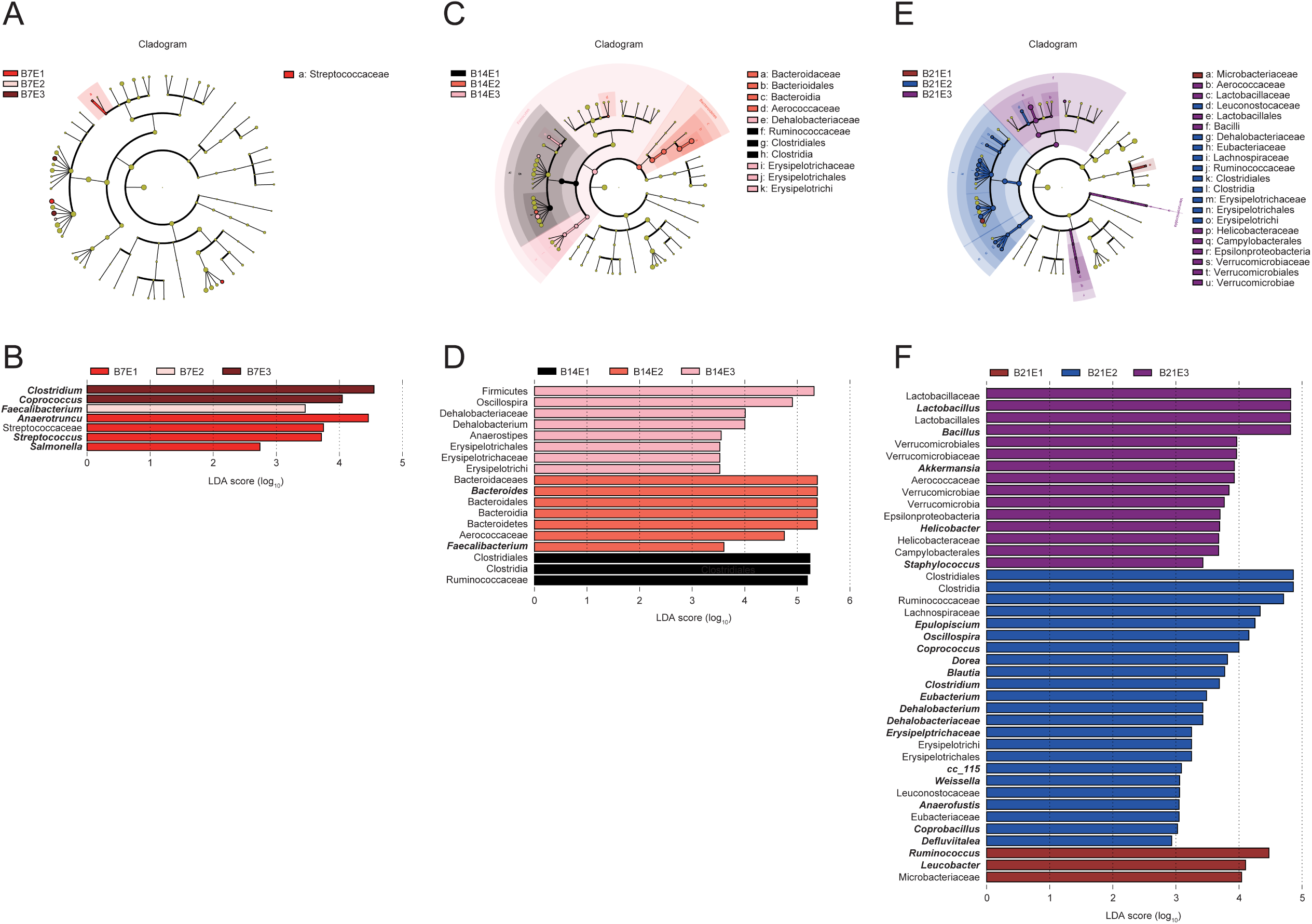
LefSe analysis of the chicken cecum microbial community in groups E1, E2, and E3 at the three sampling points. LefSe plots showing microbial strains with significant differences in groups E1, E2, and E3 (A, C, E). The different groups are represented by different colors, the microbiota that plays an important role in the different groups is represented by nodes of corresponding colors, and the organism markers are indicated by colored circles. The microbiota that does not play a significant role in the different groups is indicated by yellow nodes. From inside to outside, the circles are ordered by species at the level of phylum, class, order, family, and genus. Linear discriminant analysis (LDA) diagram (B, D, F). The different colors represent microbial groups that play a significant role in groups E1, E2, and E3. Biomarkers with statistical differences are emphasized, with the colors of the histograms representing the respective groups and the lengths representing the LDA score, which is the magnitude of the effects of significantly different species between groups.

### Metabolomic profiling of the cecum

Next, we evaluated the effects of *Salmonella* infection on the cecum metabolism of chickens on day 14 after administering enrofloxacin using PLS-DA (Figure S4). The screening conditions for the differential metabolites were as follows: (1) wariable importance in projection (VIP) ≥ 1 of the first two principal components of the PLS-DA model; (2) FC ≥ 1.2 or FC ≤ 0.83; and (3) *q* < 0.05. As shown in Table 1, in the positive ion mode, 486 differential metabolites were detected in groups E1 and E2 (219 increased, 267 decreased). Moreover, 1,577 differential metabolites were detected in groups E1 and E3 (605 increased, 972 decreased). In groups E2 and E2, 554 differential metabolites were detected (231 increased, 323 decreased). In the negative ion mode, 271 differential metabolites were detected in groups E1 and E2 (149 increased, 122 decreased). In total, 656 differential metabolites were detected in groups E1 and E3 (252 increased, 404 decreased). In total, 328 differential metabolites were detected in groups E2 and E3 (102 increased, 226 decreased; Figure 8).

**Figure 8.**
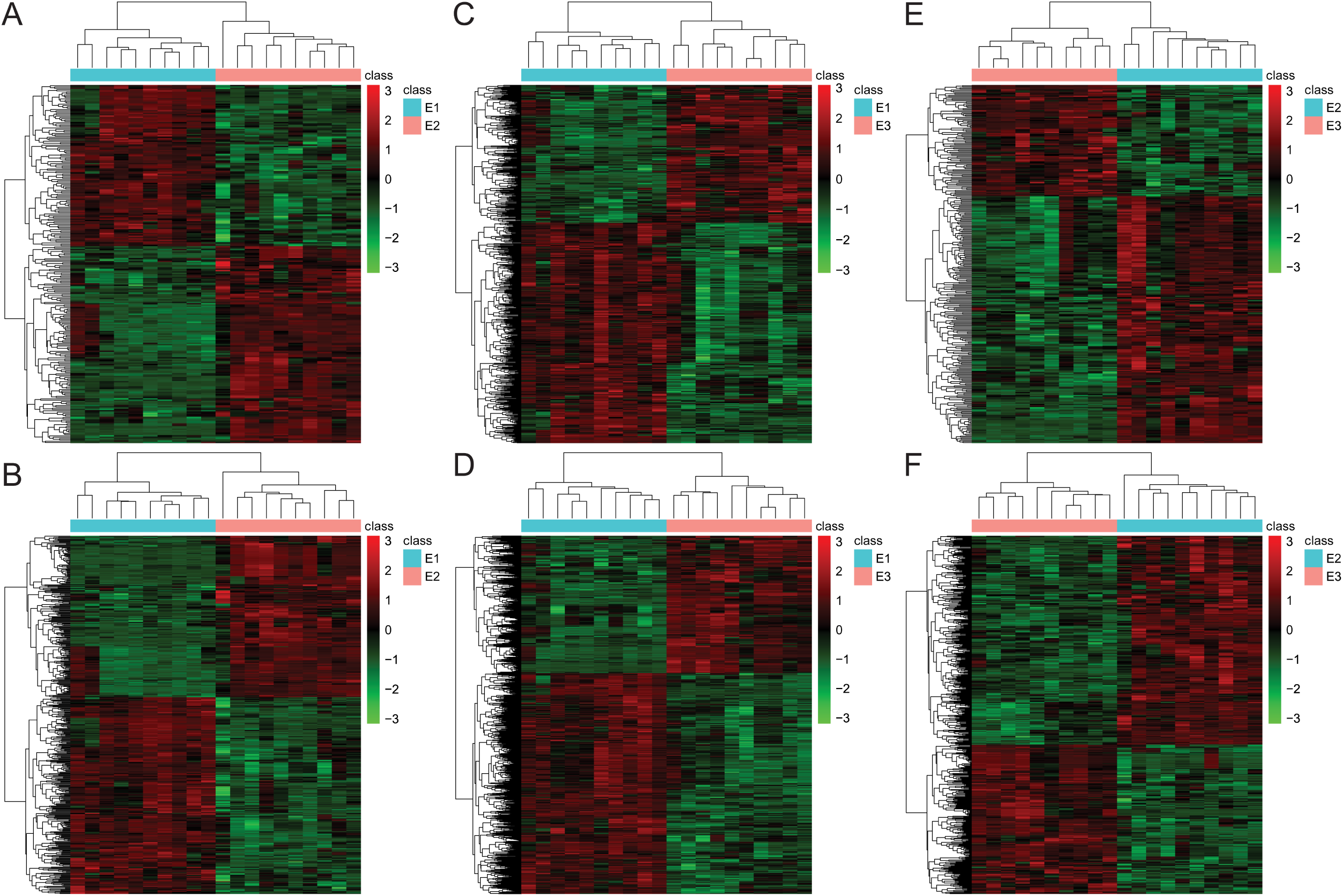
Heat map showing differential metabolites with VIP ≥ 1 for the first two principal components of the PLS-DA model, fold change (FC) ≥ 1.2 or FC ≤ 0.83, and *q* < 0.05. The green color represents downregulated differential metabolites, and the red color represents upregulated differential metabolites. (A, B) Differential metabolites in the negative or positive ion mode of group E2 versus E1; (C, D) differential metabolites in the negative or positive ion mode of group E3 versus E1; (E, F) differential metabolites in the negative or positive ion mode of group E3 versus E2.

**Table 1.**
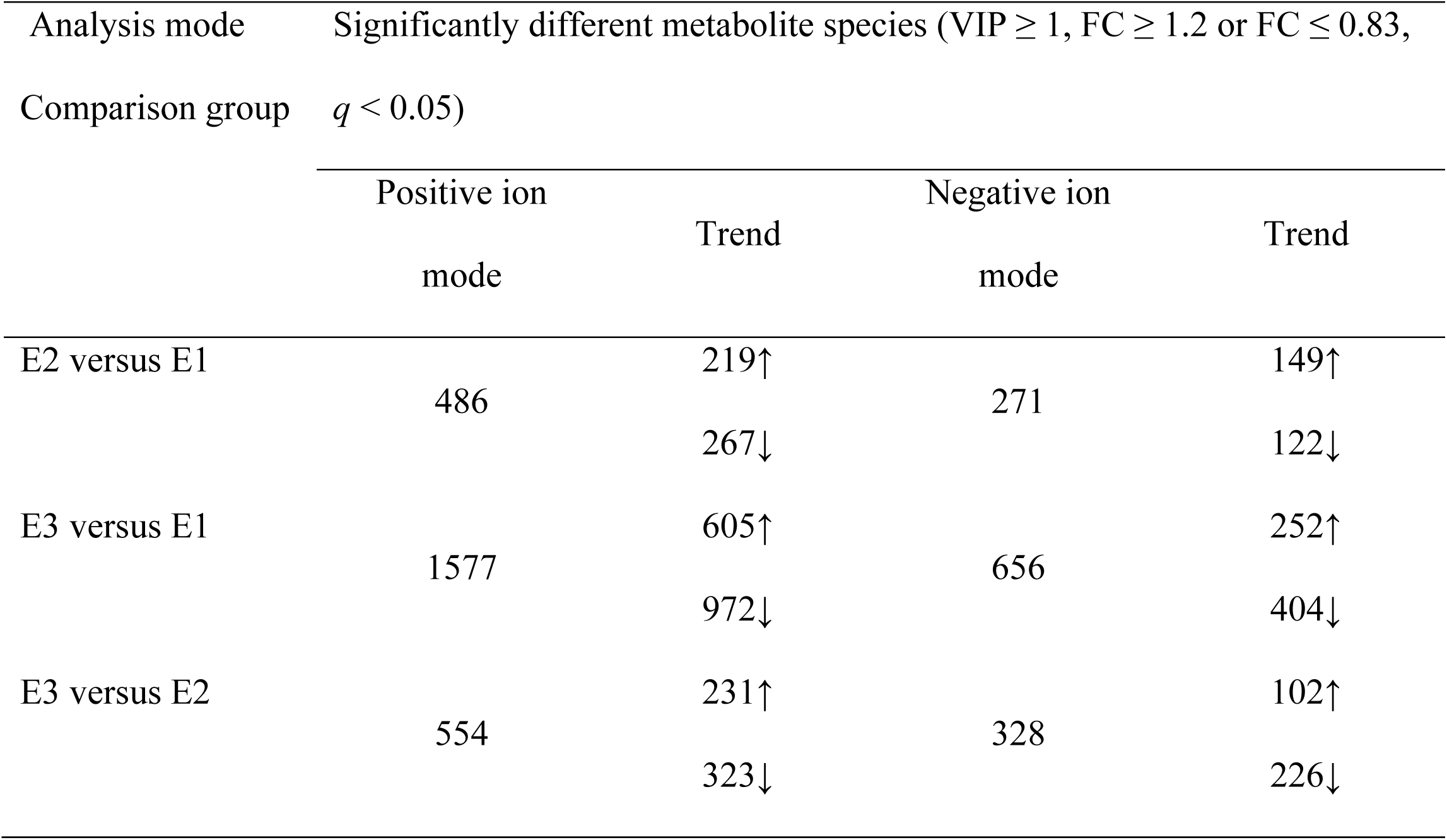
Differential metabolite statistics.

Venn diagram analysis showed that in the positive ion mode, four differential metabolites were common among the three groups, 18 differential metabolites were specific to group E2 versus E1, 437 differential metabolites were specific to group E3 versus E1, and 52 differential metabolites were specific to group E3 versus E2. In the negative ion mode, one differential metabolite was common among the three groups, 16 differential metabolites were specific to group E2 versus E1, 139 differential metabolites were specific to group E3 versus E1, and 37 differential metabolites were specific to group E3 versus E2. These results suggested that the enrofloxacin concentration affected the metabolic groups of chicken manure contents (Figure 9).

**Figure 9.**
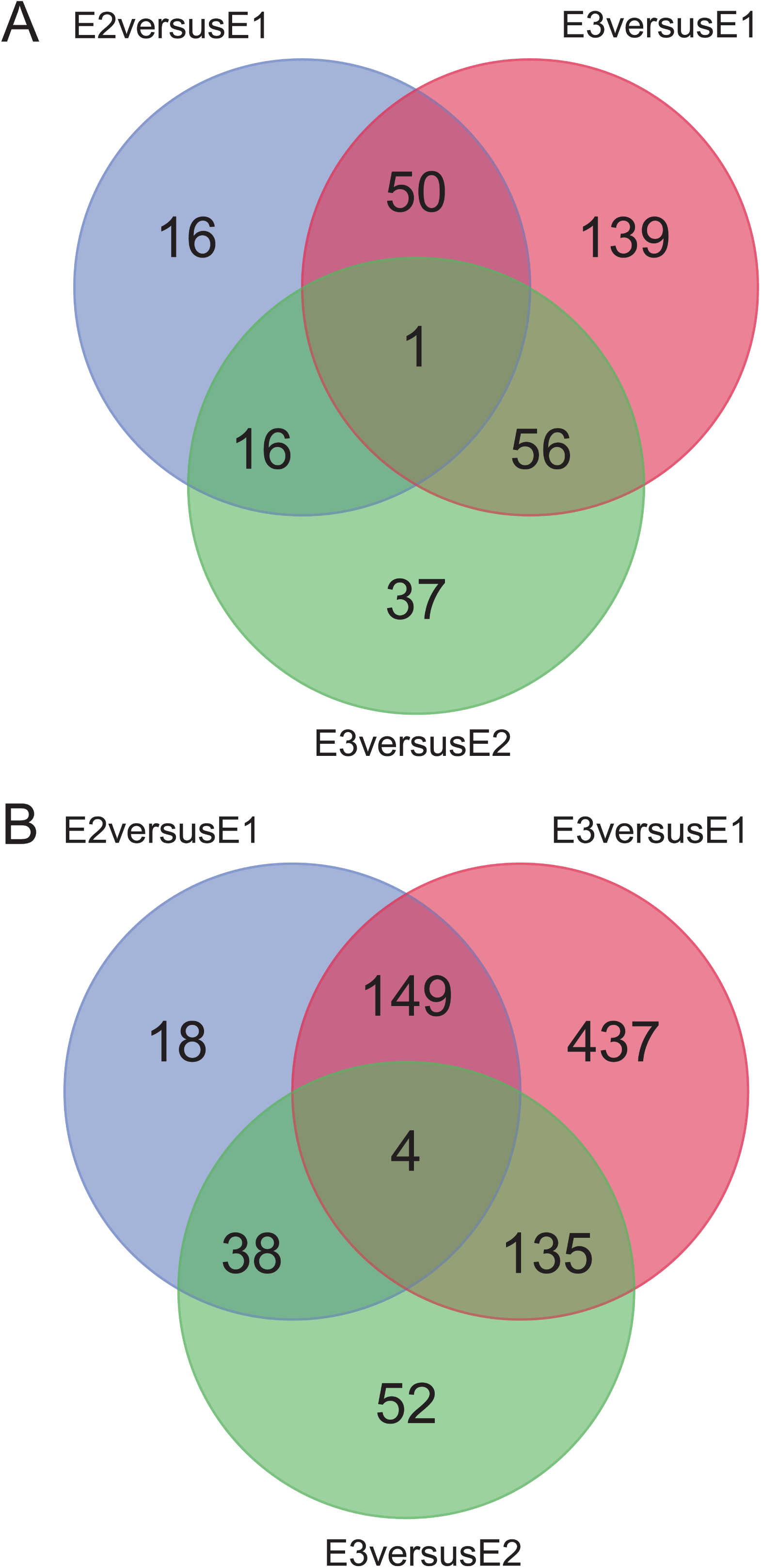
Venn diagrams showing common or unique differences in metabolites between the three groups. (A) Negative ion mode; (B) positive ion mode.

According to KEGG database analysis, in the negative ion mode, the differential metabolites of group E2 versus E1 were enriched in four metabolic pathways: vascular smooth muscle contraction (one differential metabolite), metabolic pathways (five differential metabolites), vitamin B_6_ metabolism (one differential metabolite), and lysine degradation (one differential metabolite; Figure 10A). In the positive ion mode, the differential metabolites of group E2 versus E1 were enriched in seven metabolic pathways: phenylalanine metabolism (three differential metabolites), metabolic pathways (12 differential metabolites), linoleic acid metabolism (two differential metabolites), α-linolenic acid metabolism (two differential metabolites), peroxisome proliferator-activated receptor (PPAR) signaling pathway (one differential metabolite), vascular smooth muscle contraction (one differential metabolite), and biosynthesis of amino acids (two differential metabolites; Figure 10B).

**Figure 10.**
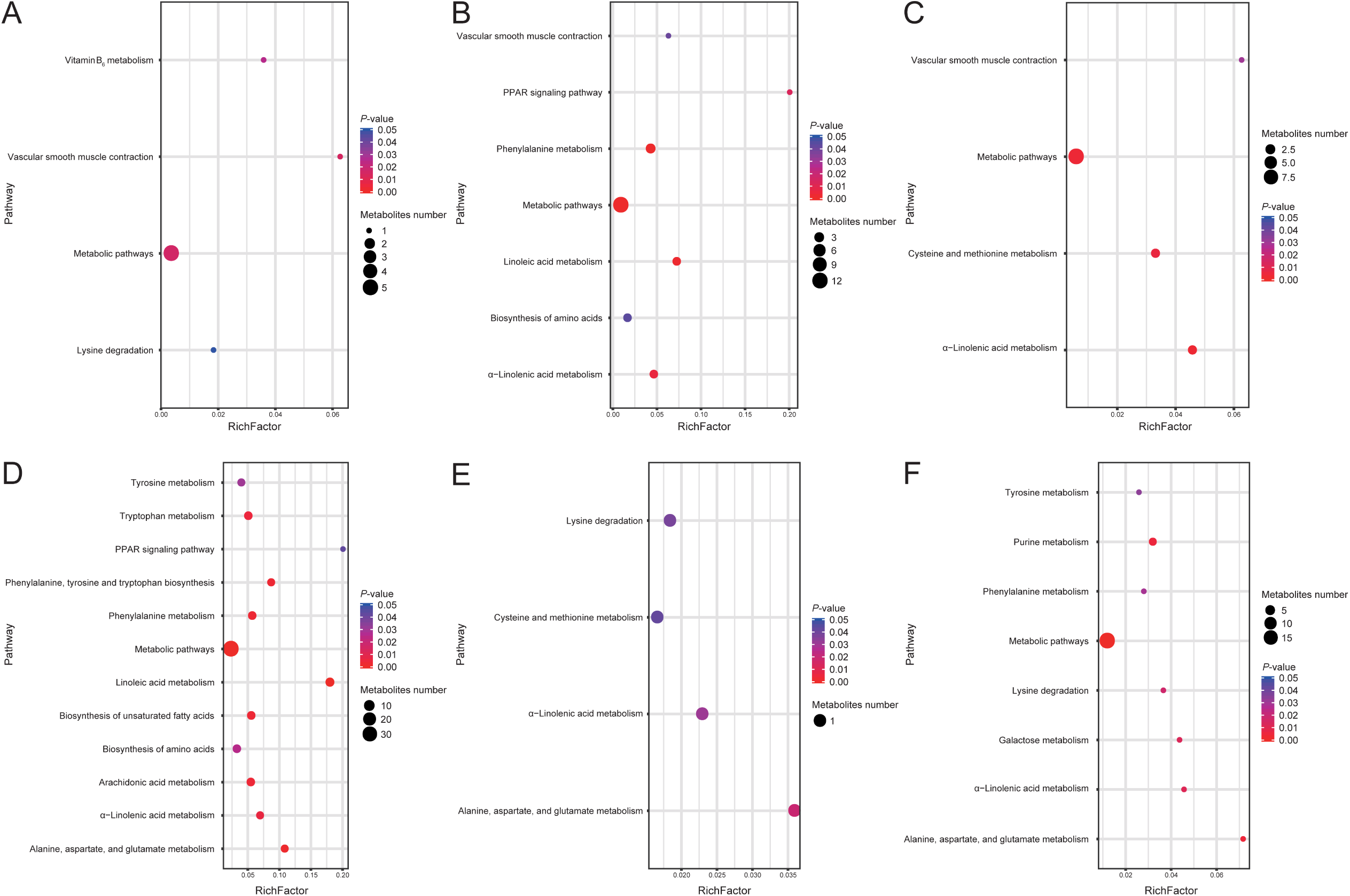
KEGG pathway analysis of metabolites in the contents of the appendix on day 14. The metabolites in the different comparison groups were analyzed for metabolic pathway enrichment on the basis of the KEGG database. The *x*-axis (RichFactor) represents the ratio of the number of differential metabolites annotated to the pathway to all the metabolites annotated to the pathway. The size of the dots represents the number of differential metabolites annotated to the pathway. (A) Upregulated metabolites in group E2 versus E1, (B) downregulated metabolites in group E2 versus E1, (C) upregulated metabolites in group E3 versus E1, (D) downregulated metabolites in group E3 versus E1, (E) upregulated metabolites in group E3 versus E2, and (F) downregulated metabolites in group E3 versus E2.

In the negative ion mode, the differential metabolites of group E3 versus E1 were enriched in four metabolic pathways: α-linolenic acid metabolism (two differential metabolites), metabolic pathways (nine differential metabolites), cysteine and methionine metabolism (two differential metabolites), and vascular smooth muscle contraction (one differential metabolite; Figure 10C). In the positive ion mode, the differential metabolites of group E3 versus E1 were enriched in 12 metabolic pathways: metabolic pathways (35 differential metabolites); linoleic acid metabolism (five differential metabolites); alanine, aspartate, and glutamate metabolism (three differential metabolites); phenylalanine, tyrosine, and tryptophan biosynthesis (three differential metabolites); phenylalanine metabolism (four differential metabolites); unsaturated fatty acids biosynthesis (four differential metabolites); arachidonic acid metabolism (four differential metabolites); tryptophan metabolism (four differential metabolites); α-linolenic acid metabolism (three differential metabolites); biosynthesis of amino acids (four differential metabolites), tyrosine metabolism (three differential metabolites), and PPAR signaling pathway (one differential metabolite; Figure 10D).

In the negative ion mode, the differential metabolites of group E3 versus E2 were enriched in four metabolic pathways: alanine, aspartate, and glutamate metabolism (one differential metabolite); α-linolenic acid metabolism (one differential metabolite); lysine degradation (one differential metabolite); and cysteine and methionine metabolism (one differential metabolite; Figure 10E). In the positive ion mode, the differential metabolites of group E3 versus E2 were enriched in eight metabolic pathways: metabolic pathways (19 differential metabolites); alanine, aspartate, and glutamate metabolism (two differential metabolites); purine metabolism (three differential metabolites); α-linolenic acid metabolism (two differential metabolites); galactose metabolism (two differential metabolites); lysine degradation (two differential metabolites); phenylalanine metabolism (two differential metabolites); and tyrosine metabolism (two differential metabolites; Figure 10F).

### Correlation analysis of the microbiome and metabolome

Next, we identified relationships among microbes and metabolites using multiomics. Bacteroidetes were negatively correlated with phosphonate and phosphinate metabolism. Verrucomicrobia were positively correlated with the intestinal immune network for IgA production, fatty acid biosynthesis, and fatty acid elongation. Firmicutes were inversely correlated with phototransduction. Cyanobacteria were positively correlated with cysteine and methionine metabolism. Tenericutes were positively correlated with glycolysis/gluconeogenesis, fatty acid metabolism, terpenoid backbone biosynthesis, fructose and mannose metabolism, and purine metabolism and negatively correlated with primary bile acid biosynthesis. Proteobacteria were positively correlated with galactose metabolism, pentose and glucuronate interconversion, arginine biosynthesis, and primary bile acid biosynthesis. Actinobacteria were negatively correlated with vitamin B_6_ metabolism and glycolysis/gluconeogenesis and positively correlated with primary bile acid biosynthesis and drug metabolism (Figure 11).

**Figure 11.**
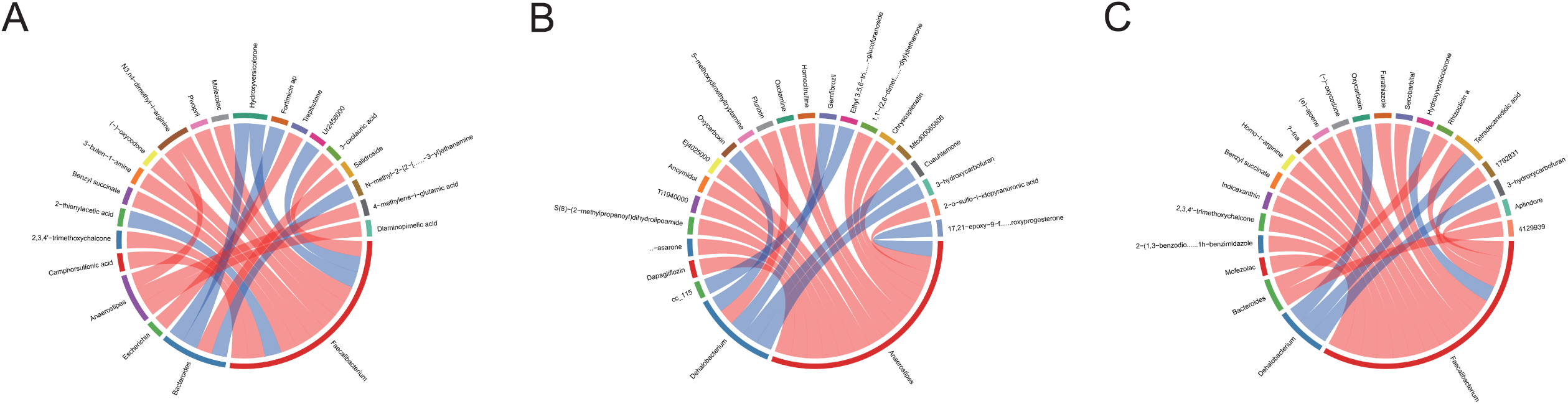
Rank correlation analysis of metabolic pathways and phylum-level microbial groups. The top 20 relationship pairs with the strongest rank correlation (with the largest correlation coefficient absolute value and a *P* value of < 0.05) are shown as chord diagrams. The blue line in the middle represents a positive correlation, and the red line represents a negative correlation.

At the genus level, rank correlation analysis was performed on differential metabolites and microbial groups. In group E2 versus E1, *Faecalibacterium* were positively correlated with three metabolites and negatively correlated with nine other metabolites, *Bacteroides* were positively correlated with three metabolites and negatively correlated with one metabolite, *Escherichia* were negatively correlated with one metabolite, and *Anaerostipes* were negatively correlated with three metabolites. In group E3 versus E1, *Dehalobacterium* were positively correlated with four metabolites and negatively correlated with one metabolite, whereas *Anaerostipes* were negatively correlated with 13 metabolites and positively correlated with one metabolite. In group E3 versus E2, *Bacteroides* were negatively correlated with two metabolites, *Dehalobacterium* were positively correlated with three metabolites, and *Faecalibacterium* were negatively correlated with 14 metabolites and positively correlated with one metabolite (Figure 12).

**Figure 12.**
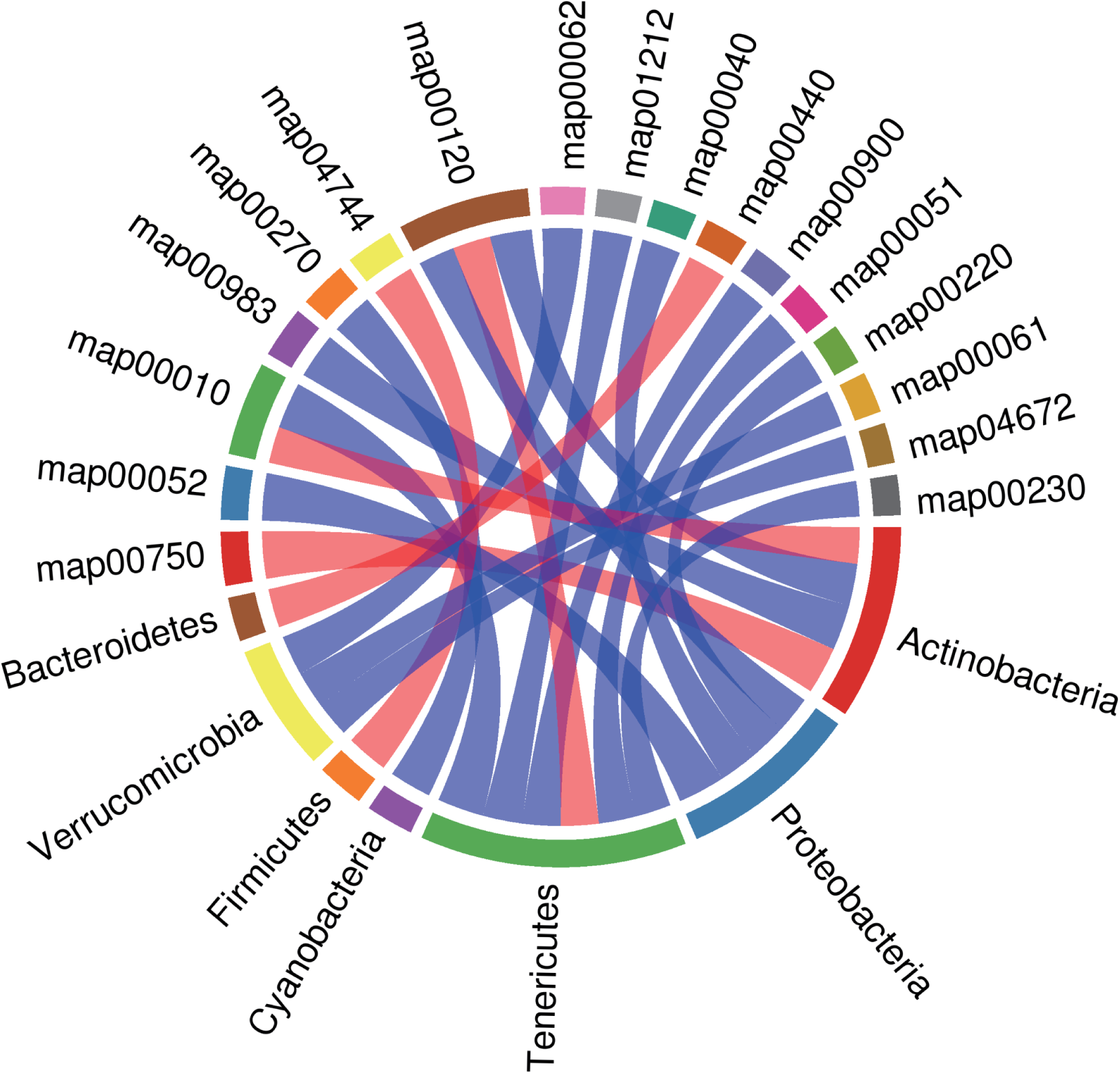
Rank correlation analysis was performed on differential metabolites and genus-level microbial groups. The top 20 relationship pairs with the strongest rank correlation (with the largest correlation coefficient absolute value and a *P* value of < 0.05) are shown as chord diagrams. The blue line in the middle represents a positive correlation, and the red line represents a negative correlation. (A) Group E2 versus E1, (B) group E3 versus E1, and (C) group E3 versus E2.

## DISCUSSION

The gut microbiome plays vital roles in resistance to exogenous microorganisms via phage deployment, secretion of antibacterial substances and competing nutrients, and intestinal barrier function. Using antibiotics destroys the gut microbiome, making the conditions suitable for pathogenic microorganisms to colonize the intestine, and affects the metabolomics. Here, we investigated the effects of enrofloxacin on the colonization of *S. Typhimurium* in the intestinal tracts of chickens. Our results showed that on days 7 and 14, *S. Typhimurium* was highly abundant in the cecum contents of mice treated with antibiotics, similar to previous reports (9,15,17). Thus, our findings showed that different concentrations of enrofloxacin altered the ability of *S. Typhimurium* to colonize the gut in chickens.

In this study, early enrofloxacin treatment suppressed *S. Typhimurium* colonization. However, as the gut microbial community matured, the abundance of *S. Typhimurium* in untreated chickens was gradually decreased, whereas that in enrofloxacin-treated chickens gradually increased and persisted. These results were similar to those of a previous study (15) and suggested that enrofloxacin may cause severe *S. Typhimurium* infection.

On day 7, the highest diversity was observed in the high-dose group, and the lowest diversity was observed in the low-dose group. On day 21, the low-dose group showed the highest diversity, whereas the high-dose group showed the lowest diversity. These results are different from previous findings, potentially because of the use of antibiotics to challenge *Salmonella* in this study (9,18,19). Generally, the alpha diversity of the cecum flora increases with age. A higher flora diversity is thought to be a marker of mature intestinal flora that is less sensitive to environmental factors and less susceptible to infection (20). Previous studies have shown that mice with lower intestinal flora complexity are less resistant to *Streptococcus enterocolitis* colonization and more susceptible to enterocolitis than normal mice (21).

Our findings showed that using antibiotics significantly altered the microbial community structure, with clear differences according to the antibiotic concentration. The main reasons for the changes in microbial structure were decreased Proteobacteria and Firmicutes and increased Bacteroidetes at the three sampling points. Indeed, the early microbial community mainly comprised Firmicutes and Proteobacteria, accounting for more than 90% of the total sequence, whereas the late microbial community was dominated by Bacteroidetes and Firmicutes. Studies have shown that Bacteroides strains can produce propionate to reduce the colonization of *S. Typhimurium* (10).

Firmicutes are primarily anaerobic bacteria. The anatomic physiology and feeding habits of chickens may explain the high abundance of Firmicutes. Turnbaugh et al. found that the appendices of obese mice contain higher concentrations of acetic acid and butyric acid and that the ratio of Firmicutes to Bacteroidetes is increased in comparison with normal mice (22). Changes in the ratio of Firmicutes to Bacteroidetes in the experimental group may affect the accumulation of chicken fat and the production of SCFAs, which may explain the higher abundance of *S. Typhimurium* in group E2 on day 14. Additionally, on day 14, the abundances of Enterobacteriaceae in groups E2 and E3 were lower than that in group E1, although the difference was not significant.

Enterobacteriaceae can inhibit the colonization of *Salmonella* in the intestine by competing for oxygen (13). Notably, the abundances of *Anaerotruncus*, *Butyricicoccus*, *Ruminococcus*, *Dorea*, and *Clostridium* were gradually decreased. In addition, higher concentrations of enrofloxacin resulted in lower abundances of *Ruminococcus* and *Anaerotruncus* on days 7 and 14, similar to a previous study of florfenicol (23).

Many anaerobic bacteria can produce SCFAs via microbial fermentation of carbohydrates, which have immunomodulatory and anti-inflammatory effects. Butyric acid can mediate the inhibition of *Salmonella* by downregulating the expression of pathogenic island I and can also reduce *Salmonella*-induced pro-inflammatory responses to intestinal cells *in vitro* (24,25). These aforementioned bacteria compete with *Salmonella* for oxygen and produce high concentrations of SCFAs to limit their colonization (26). SCFAs exert inflammatory effects by regulating the production of cytokines and prostaglandins (27) and can enhance host immune function. Sunkara et al. found that SCFAs reduce the *Salmonella* load in the intestinal contents and enhance the host’s defense ability against *Salmonella* (28). Butyrate can promote the β-oxidation of colonic epithelial cells, reduce the oxygen partial pressure in the intestinal cavity, and inhibit pathogenic bacteria (29). Any reduction in SCFA production could explain the high abundance of *Salmonella* in the cecum of antibiotic-treated chickens. In this study, on day 14, *Lactobacillus* and *Oscillospira* numbers were higher in groups E2 and E3 than in group E1, consistent with previous studies. This may be due to the microaerophilic growth of *Lactobacillus* and the production of reactive oxygen species by granulocytes in the infected chicken cecum to penetrate the site of inflammation and provide facorable growth conditions for lactic acid bacteria (26,30,31). Notably, accumulation of lactic acid may damage the intestinal defense barrier and increase the osmotic load in the intestinal cavity, and *Salmonella* can oxidize lactic acid to enhance their intestinal adaptability (14,32). Thus, our results indicated that enrofloxacin decreased the colonization resistance of the original *S. Typhimurium* in the intestine, allowing *S. Typhimurium* to more easily colonize the intestine.

Tumor necrosis factor-α (*TNF-α*) and *MUC*2 genes were upregulated in the cecum of chickens infected with *S. Typhimurium* following enrofloxacin treatment. These genes are important indicators for evaluating the inflammatory response in birds and can induce an inflammatory response following pathogen infection. *MUC*2 is the most important mucin (33). *Salmonella* can use type I flagellates to bind to mannose residues distributed on the surface of the mucin layer to attach to the mucin surface and colonize the intestine (34). In this study, *IL-*18 expression was inhibited in the enrofloxacin group. IL-18 can induce Th1-type cells and NK cells to produce IFN-γ, thereby promoting neutrophil and macrophage killing and clearance of intracellular bacteria. In addition, downregulation of IL-18 increases bacterial load and decreases survival in the liver and spleen (35,36).

Occludin (OCLN) is a major tight junction protein in the intestine. In this study, gene expression was significantly downregulated in group E3, suggesting that high concentrations of enrofloxacin may diffuse more effectively throughout the body by reducing tight junction activity between the intestinal tissue and intestinal mucosal epithelial cells. Additionally, the expression of immune-related genes was significantly altered in chickens infected with *S. Typhimurium* after enrofloxacin administration, with increased pro-inflammatory gene expression and decreased anti-inflammatory gene expression in comparison with the untreated group. These findings suggested that enhanced *S. Typhimurium* colonization may cause a more severe inflammatory response in the intestinal tract of chickens.

Metabolomic analysis showed that the differential metabolic pathways shared by groups E2 versus E1 and E3 versus E1 included linoleic acid metabolism and α-linolenic acid metabolism. According to several reports, polyunsaturated fatty acids, such as linoleic acid, α-linolenic acid, and γ-linoleic acid, are metabolized by macrophages into fatty acids with a ketone structure. Notably, fatty acids containing an enone structure have the most effective anti-inflammatory activity (37).

Linolenic acid can kill *Helicobacter pylori* and reduce the bacterial load in the stomachs of mice. *In vivo*, treatment with linolenic acid reduces the levels of pro-inflammatory cytokines (e.g., IL-1β, IL-6, and TNF-α) (38). Interestingly, addition of flaxseed oil to the diet of pigs recovered the levels of α-linolenic acid, eicosapentaenoic acid, and total n-3 polyunsaturated fatty acids in the intestinal tract and improved intestinal morphology. Flaxseed oil can also increase jejunal lactase activity and claudin-1 protein expression; downregulate the mRNA expression of intestinal necrosis signals; downregulate intestinal *TLR*4 mRNA, its downstream signals (*MyD*88, *NF-κB*, *NOD*1, and *NOD*2), and their adaptor molecules; and decrease the functionality of RIPK2 expression (39).

Notably, the changes observed in the PPAR signaling pathway and amino acid biosynthesis were due to the metabolic effects of *S. Typhimurium* colonization in the chicken cecum. The abundance of α-linolenic acid was lower in groups E2 and E3 than in group E1 and lower in group E3 than in group E2. This decrease in the level of α-linolenic acid may be the reason for the high abundance of *S. Typhimurium* in groups E2 and E3. However, further studies are required to elucidate the effects of administering α-linolenic acid on *S. Typhimurium* colonization.

In this study, we found that IgA production by the *Verrucomicrobia* intestinal immune network was positively related to fatty acid biosynthesis. The genus *Verrucomicrobia* contains only one member, i.e., *Akkermansia muciniphila*, which is a member of the human intestinal flora and a common bacterium that degrade mucins in the gut (40,41). Colonization of *Aspergillus* slime has been reported to be protective against diet-induced obesity (42,43) and to promote mucosal wound healing (44) and antitumor responses during anti-programmed death-1 immunotherapy (45). Therefore, when enrofloxacin is administered, changes in *Verrucomicrobia* may affect *S. Typhimurium* colonization.

In this study, we found that *Faecalibacterium* was associated with the most metabolites in the E2 versus E1 and E3 versus E2 groups and that *Anaerostipes* was associated with the most metabolites in the E3 versus E1 group. Several studies have reported that the metabolites secreted by *Faecalibacterium prausnitzii* attenuate inflammation in cells and 2,4,6-trinitrobenzenesulphonic acid-induced colitis models by blocking nuclear factor (NF)-κB activation and IL-8 production (46). Notably, *F. prausnitzii* secretes a protein with a molecular weight of 15 kDa, which mediates resistance to colitis in animal models by inhibiting the NF-κB pathway in intestinal epithelial cells (47). Moreover, *F. prausnitzii* can adapt to the mucus layer and ferment to produce butyrate. The absence of *F. prausnitzii* and other microorganisms with similar functions is often accompanied by inflammatory bowel disease and obesity (48). These bacteria may be the major bacteria affecting the intestinal metabolome following enrofloxacin administration.

Our results indicated that infection with *S. Typhimurium* after enrofloxacin administration affected the colony structure, metabolite composition, and cecum gene expression in the chicken cecum, thereby altering *S. Typhimurium* colonization in the gut. The effects varied according to the concentration of enrofloxacin used. The colonies and metabolites identified in this study may provide important insights into the prevention and control of *S. Typhimurium* infection. The observed increase in *S. Typhimurium* infection induced by enrofloxacin also facilitate the more careful and rational use of antibiotics in poultry.

## MATERIALS AND METHODS

### Bacterial strain

*S. Typhimurium* ATCC14028 (stored in our laboratory) was incubated in lysogeny broth at 37°C for 16 h. For bacterial counting, the liquid was diluted and inoculated on XLT4 agar plates at 37°C for 24 h.

### Enrofloxacin exposure and infection in chickens

The experimental protocol involving animals in this study was approved by the Animal Ethics Committee of Sichuan University. One-day-old specific-pathogen-free chickens (n = 180) were randomly divided into six groups (n = 30 each). Each group was housed in a separate isolator containing drug-free feed and water. From 1 to 5 days of age, chickens were fed different concentrations for enrofloxacin (C1 and E1 [control group], 200 μL distilled water; C2 and E2, 200 μL of 10 mg/kg enrofloxacin; C3 and E3, 200 μL of 100 mg/kg enrofloxacin). At 6 days of age, chickens in groups E1, E2, and E3 were challenged with oral administration of 1 × 10^8^ colony-forming units (CFU) *S. Typhimurium*, whereas chickens in groups C1, C2, and C3 were given equal amounts of Luria-Bertani broth. All chickens were euthanized on days 7, 14, or 21 (n = 10/group at each time), and cecum contents, heart, spleen, liver, and ceca were collected. All cecum contents were collected from the same part of the cecum. The ileum and colon were rapidly clamped to avoid overflow of gastrointestinal digestive fluids, which can contaminate other parts of the intestine (Figure 13).

**Figure 13.**
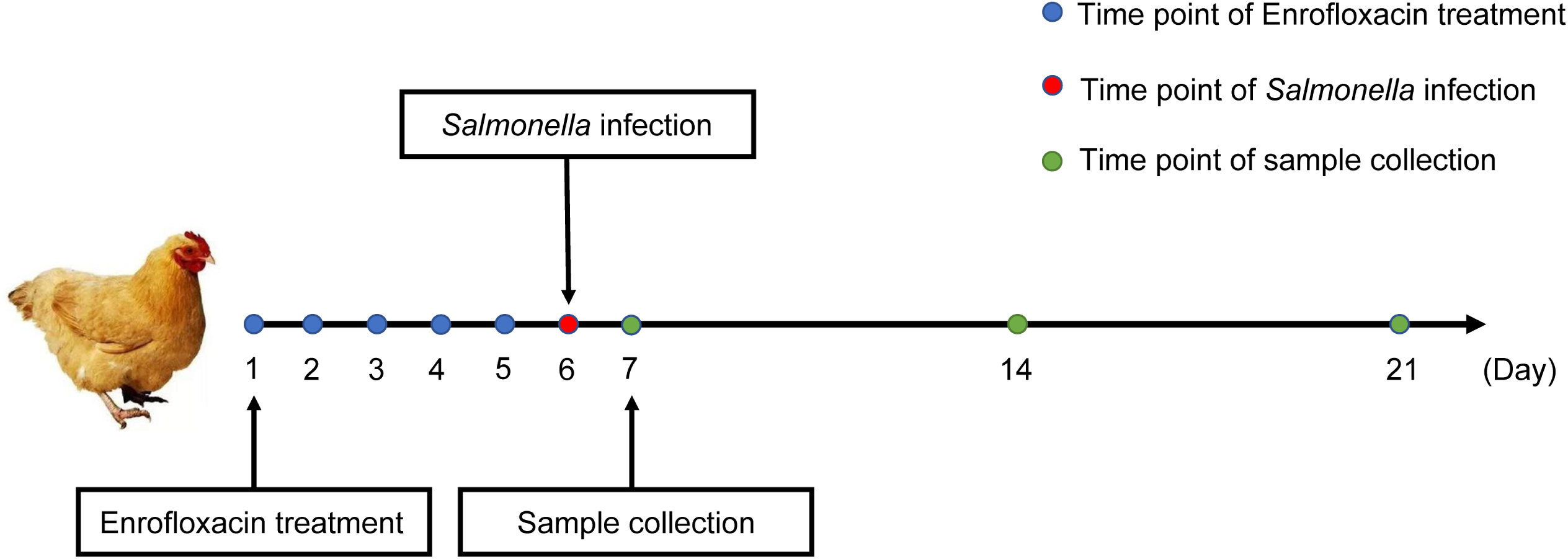
Animal experiment design. In total, 180 specific-pathogen-free chickens were randomly divided into six groups (C1, C2, C3, E1, E2, and E3). Groups E1, E2, and E3 were fed *Salmonella* Pullorum 1 × 10^8^ colony-forming units (CFUs)/mL per chicken at 6 days of age, and groups E1 and C1 were fed equal amounts of lactose broth. Samples were taken from each chicken at 7, 14, and 21 days of age.

### Total RNA extraction

Total RNA from chicken cecum tissues previously collected on day 14 was extracted using TRIzol reagent (Invitrogen, Carlsbad, CA, USA) according to the manufacturer’s instructions. Approximately 60 mg tissue was ground into a powder in the presence of liquid nitrogen in a 2-mL tube, followed by homogenization for 2 min and then resting in a horizontal position for 5 min. The mixture was centrifuged (12,000 × *g*) at 4°C for 5 min, and the supernatant was then taken and placed in a new Eppendorf tube. A mixture of 0.3 mL chloroform/isoamyl alcohol (24:1) was injected, shaken vigorously for 15 s, and then centrifuged at 4°C for 10 min at 12,000 × *g*. The remaining upper aqueous phase RNA was then transferred to a new tube, and an equal volume of isopropanol alcohol was added, followed by centrifugation at 4°C for 20 min at 12,000 × *g*. After discarding the supernatant, the RNA pellet was washed twice with 1 mL of 75% ethanol, and the mixture was centrifuged at 12,000 × *g* for 3 min at 4°C to collect any residual ethanol. The pellet was air dried for 5–10 min in a biosafety cabinet, and RNA was dissolved with 25–100 μL diethylpyrocarbonate-treated water. Subsequently, a spectrophotometer and bioanalyzer were used for total RNA characterization and quantification.

### 16S rRNA amplicon sequencing

Samples were collected from 10 chickens from the same group at each sampling time. According to the manufacturer’s instructions, a QIAamp DNA Stool Mini Kit (Qiagen, Hilden, Germany) was used to test 200 mg aliquots and 30 mg tissue from each chicken manure sample. Total genomic DNA was extracted from the samples, and the concentration and quality of the extracted DNA were measured using a NanoDrop spectrophotometer and agarose, followed by confirmation using gel electrophoresis. The extracted DNA was stored at −20°C until further analysis.

Using the isolated genomic DNA as the template, polymerase chain reaction (PCR) amplification was performed on the V3-V4 hypervariable region of the bacterial 16S rRNA gene. The primers used were 338F (5′-ACTCCTACGGGAGGCAGCAG-3′) and 806R (5′-GGACTACHVGGGTWTCTAAT-3′). Sample-specific 7-bp barcodes were assigned to multiplex sequencing primers. The 25-μL PCR system and PCR reaction parameters for PCR amplification were set up using Agencourt. AMPure XP magnetic beads were used for purification of PCR amplification products, and products were dissolved in elution buffer. The libraries were labeled, and the fragment range and concentration of the libraries were checked using an Agilent 2100 Bioanalyzer. The qualified libraries were sequenced on a HiSeq platform according to the inserted fragment size.

### 16S rRNA sequencing data analysis

Both off-machine data filtering and the remaining high-quality clean data are used for analyses. Overlaps between reads were used to stitch the reads into tags, cluster the tags into operational taxonomic units (OTUs), and compare them with the database. The reads were also used for species annotation. Sample species complexity analysis and analysis of species differences between groups were performed according to the OTU and annotation results as well as correlation analysis and model prediction. The original sequencing data were processed as follows to obtain clean data. For sequence splicing, Fast Length Adjustment of SHort reads software (FLASH, v.1.2.11) (49) was used to assemble the paired reads obtained by paired-end sequencing into a sequence using overlapping relationships to obtain tags in hypervariable regions.

Generally, OTUs are a unified flag set for taxonomic units (strain, genus, species, grouping, etc.), facilitating analyses in phylogenetic or population genetic studies. To elucidate the bacterial number, genus, and other information in the sequencing results of a sample, clustering operations were performed. Using the classification operation, sequences were classified into many groups according to their similarities, with each group being considered an OTU. Tags with a similarity above 97% were clustered into an OTU.

After obtaining the OTU representative sequence, RDP Classifier software (v.2.2) (50) was used to compare this sequence with the database for species annotation (confidence threshold: 0.8). To obtain the species classification information corresponding to each OTU, the RDP Classifier’s Bayesian algorithm was used to conduct a taxonomic analysis on the OTU representative sequences and for statistical analysis of the phylum, class, order, family, and genus level composition for each sample.

### Bioinformatics and statistical analysis

Alpha diversity is an analysis of the diversity of species in a single sample. The richer the species in the sample, the larger the observed species index, Chao index, ACE index, Shannon index, and Simpson index and the smaller the Good coverage index (51). In addition, higher Good coverage values indicated lower the probability that the sequence in the sample was not detected. Beta-diversity analyses (51-53) were used to compare differences between pairs of samples in terms of species diversity. In this analysis, the content of each species in the sample was first analyzed, and the beta-diversity value was then calculated between samples. Beta-diversity analysis was performed using QIIME (54) (v.1.80). Linear discriminant analysis (LDA) effect size (LefSe) (55) was used to identify high-dimensional biomarkers and reveal genomic features based on genes, metabolism, and classifications for distinguishing between two or more biological taxa.

### Untargeted metabolomics for chicken cecal content

After thawing samples slowly at 4°C, 25 mg was measured and added to a 1.5-mL Eppendorf tube. Then, 800 μL extract (methanol/acetonitrile/water, 2:2:1, v/v/v, precooled at −20°C), 10 μL internal standard 1, and 10 μL internal standard 2 were added. Two small steel balls were then added, and samples were ground in a tissue grinder (50 Hz, 5 min), sonicated at 4°C for 10 min, and placed in a freezer at −20°C for 1 h. After centrifugation, 600 μL supernatant was removed, placed in a refrigerated vacuum concentrator, and dried. Next, 200 μL of a double solution (methanol/H_2_O, 1:9, v/v) was added for recombination, and the samples were vortexed for 1 min and sonicated for 10 min at 4°C. The supernatants were then centrifuged at 25,000 × *g* for 15 min at 4°C and placed in a sample vial. Then, 20 μL supernatant from each sample was mixed with quality control samples to assess the repeatability and stability of the liquid chromatography-mass spectrometry (LC-MS) analysis. A BEH C18 column (1.7 μm × 2.1 mm × 100 mm; Waters, Milford, MA, USA) was used for metabolite isolation. The mobile phases used in the positive ion mode were an aqueous solution containing 0.1% formic acid (liquid A) and a solution of 100% methanol containing 0.1% formic acid (liquid B). The mobile phases in the negative ion mode were an aqueous solution containing 10 mM ammonium formate (liquid A) and a solution of 10 mM ammonium formate and 95% methanol (liquid B). A Q Exactive mass spectrometer (Thermo Fisher Scientific) was used for primary and secondary mass spectrometry data acquisition. All LC-MS/MS mass spectrometry raw data (original files) were sent to Compound Discoverer (v.3.0; Thermo Fisher Scientific) for data processing. Identification of metabolites was performed using several databases, such as the BGI Library, mzCloud, and ChemSpider (HMDB, KEGG, and LipidMAPS). Results from Compound Discoverer v.3.0 were imported into metaX (56) for data preprocessing. All the identified metabolites were classified and annotated with reference to the Kyoto Encyclopedia of Genes and Genomes (KEGG) and HMDB databases to understand the classification of the metabolites. Multivariate statistical analysis (principal component analysis [PCA] and partial least squares discriminant analysis [PLS-DA]) and univariate analysis (fold change [FC] and Student’s *t*-tests) were used to screen for between-group differential metabolites. Here, PCA and PLS-DA (57,58) were used to model the relationship between metabolite expression and sample category, which allowed the prediction of the sample category, combined with the multiplier of difference and *t*-test, to finally identify the differences in metabolites between groups. Data were analyzed using clustering analysis, log_2_ conversion, and *z*-score normalization (zero mean normalization). Hierarchical clustering was used in the clustering algorithm, and the Euclidean distance was used to calculate the metabolic pathway enrichment of differential metabolites in the KEGG database.

### Bacterial enumeration

First, 200 mg organ tissue and cecum content from each animal was added to 10 mL sterile saline and shaken well. Then, 1 mL bacterial solution was added to 9 mL sterile saline for homogenization; samples were serially diluted and spread on XLT4 agar plates. The medium was then incubated for 24 h under aerobic conditions at 37°C, and *S. Typhimurium* colonies were counted. The results are expressed as log_10_ CFU/g. All analyses were performed in triplicate.

### Real-time quantitative reverse transcription PCR (qPCR)

Total RNA was extracted as previously described. The quality and concentration of RNA were determined using a Nanodrop 2000 spectrophotometer. Briefly, 1 μg total RNA from each sample was reverse-transcribed into cDNA using SuperScript II (Life Technologies, Invitrogen). The primers used for reverse transcription were oligonucleotide (dT) primers and random hexamers. Quantitative PCR was performed using 2 × T5 Fast qPCR Mix (SYBR Green I) with a Bio-Rad CFX Real-Time PCR Detection System (Bio-Rad Laboratories, Hercules, CA, USA) according to the manufacturer’s protocol. Table S1 shows the primers used in this study. The relative mRNA expression levels of each target gene were calculated using the 2^−ΔΔCT^ method with β-actin as the endogene. In all cases, all samples were reacted in triplicate in parallel.

### Correlation analysis between the microbiome and metabolome

Pretreatment of the collected positive and negative ions in the metabolome yielded 3,417 metabolites, which were downscaled using gene co-expression network analysis, yielding 42 co-expression clusters. In total, 424 of the 3,417 metabolites were annotated into 95 pathways, and pathway abundance was calculated on the basis of metabolite intensity. Correlations between metabolite intensity and microbial abundance were assessed using Spearman’s correlation analysis.

### Statistical analysis

*S. Typhimurium* count results are expressed in log_10_ CFU/g. The relative mRNA expression levels of each immune gene were calculated using the 2^-ΔΔCT^ method. One-factor ANOVA was performed on the above results using SPASS 20.0.

## ACKNOWLEDGMENTS

This work was supported by the General Program of National Natural Science Foundation of China (grant nos. 31830098 and 3177131163), the China Agriculture Research System National System for Layer Production Technology (grant no. CARS-40-K14), the Key Research and Development Projects in Sichuan Province (grant no. 2014NZ0002), and the Fundamental Research Funds for the Central Universities (grant no. SCU2019D013).

---

**Figure S1.** Rarefaction curves for chicken-blind flora. (A) ACE index, (B) Chao1 index, (C) Good coverage index, (D) observed species index, (E) Shannon index, and (F) Simpson index.

**Figure S2.** Alpha-diversity analysis of the cecum microbiota after *Salmonella* infection. (A, B, C) Alpha diversity in groups E1, E2, and E3 on days 7, 14, and 21. (D, E, F) Alpha diversity in groups E1, E2, and E3 at the three sampling points.

**Figure S3.** Beta-diversity analysis of the cecum microbiota after *Salmonella* infection. Principal component analysis (PCA), in which the absolute abundance of operational taxonomic units (OTUs) in groups E1, E2, and E3 at the three sampling points was used to calculate the distribution of each sample. (A, B, C) PCA for groups E1, E2, and E3 on days 7, 14, and 21. (D, E, F) Cluster tree for groups E1, E2, and E3 at the three sampling points.

**Figure S4.** Partial least squares discriminant analysis (PLS-DA) fractional scatterplots of the identified metabolites with the first principal component on the horizontal axis and the second principal component on the vertical axis. The number in parentheses indicates the score for that principal component and represents the percentage of the overall variance explained by the corresponding principal component. (A, B) For metabolites in the negative or positive ion mode of group E2 versus E1; (C, D) for metabolites in the negative or positive ion mode of group E3 versus E1; (E, F) for metabolites in the negative or positive ion mode of group E3 versus E2.

